# RNA-seq based analysis of population structure within the maize inbred B73

**DOI:** 10.1101/043513

**Authors:** Zhikai Liang, James C. Schnable

## Abstract

B73 is a variety of maize (Zea mays ssp. mays) widely used in genetic, genomic, and phenotypic research around the world. B73 was also served as the reference genotype for the original maize genome sequencing project. The advent of large-scale RNA-sequencing as a method of measuring gene expression presents a unique opportunity to assess the level of relatedness among individuals identified as variety B73. The level of haplotype conservation and divergence across the genome were assessed using 27 RNA-seq data sets from 20 independent research groups in three countries. Several clearly distinct clades were identified among putatively B73 samples. A number of these blocks were defined by the presence of clearly defined genomic blocks containing a haplotype which did not match the published B73 reference genome. In a number of cases the relationship among B73 samples generated by different research groups recapitulated mentor/mentee relationships within the maize genetics community. A number of regions with distinct, dissimilar, haplotypes were identified in our study. However, when considering the age of the B73 accession – greater than 40 years – and the challenges of maintaining isogenic lines of a naturally outcrossing species, a strikingly high overall level of conservation was exhibited among B73 samples from around the globe.

## Background

A great deal of biological research depends on reference genotypes that allow researchers around the world on work with material that is genetically identical or nearly identical. For many decades, assessing whether two samples labeled as coming from genetically identical sources truly were identical was a costly, time consuming, and often inconclusive process. Recent advances in genotyping and sequencing technology have revealed a number of cases where sample names and sequence information significantly different stories. One study of human cell cultures found that 18% of cell lines were either contaminated or something entirely different from what they were labeled as [1] with the widely used HeLa cell line being one of the most frequent offenders [2]. Among plants, a recent resequencing study of arabidopsis demonstrated that a line believed to carry a mutation for the ABP1 gene in an otherwise Col-0 background actually contained a wide range of other nonsense and missense mutations as well as a large region on chromosome 3 which came from a different arabidopsis accession [3]. In soybean (Glycine max), segregating variation was observed among various inbred sources of the line Williams82 which was used in the construction of the soybean reference genome [4].

Here we set out to quantify how severely these issues of divergence among samples labeled as belonging to the same genetic background impact maize (Zea mays), the preeminent model for plant genetics over the past 100 years. Unlike soybean and arabidopsis, maize is a naturally outcrossing species, so reference genotypes must be maintained by controlled self-pollination in each generation. This study focuses specifically on the maize reference genotype B73, which was developed in Iowa and first registered in 1972 [5], widely used in commercial hybrid seed production across the United States for much of the 1970s and 1980s [6] and is represented in the parentage of many elite lines even today [7]. B73 has also been widely used by plant biologists conducting basic genetic research in maize, and was employed in the sequencing and assembly of the first maize reference genome [8].

## Methods

### Data sources

A search of NCBI’s sequence read archive identified 25 Illumina RNA-seq data sets deposited by 19 independent research group in three countries (Table 1). Two additional RNA-seq data sets were constructed from B73 seed requested from Iowa State and the USDA’s Germplasm Resources Information Network (Control 1 and Control 2 respectively). For these two samples RNA was extracted from 12-day old B73 seedlings grown at the University of Nebraska-Lincoln (Table 1). In four cases where the total amount of data per run was limited (USA 6, USA 8, USA 9 and USA 17), data from multiple sequencing runs labeled as coming from the same sample were grouped together for analysis. In one case, SRR514100, the total quantity of data was excessive, so only 1/10th of the total data set was employed.

**Table 1.** B73 RNA-seq data sets sources.

| Sample Name | Run Accession | Library Layout (bp) | Institute |
| --- | --- | --- | --- |
| Control 1 | - | Paired (101) | University of Nebraska - Lincoln |
| Control 2 | - | Single (51) | University of Nebraska - Lincoln |
| USA 1[9] | SRR651051 | Paired (51) | University of Minnesota |
| USA 2[10] | SRR1819621 | Paired (52) | University of Minnesota |
| USA 3[11] | SRR404150 | Single (76) | University of Wisconsin - Madison |
| USA 4[12] | SRR514100 | Paired (151) | University of Wisconsin - Madison |
| USA 5[13] | SRR940300 | Single (101) | University of Wisconsin - Madison |
| USA 6[14] | *SRR395191, SRR395192*<br>*SRR395194, SRR395208* | Single (40) | Iowa State University |
| USA 7 | SRR445245 | Paired (102) | Iowa State University |
| USA 8[15] | *SRR039505, SRR039506* | Single (35) | Danold Danforth Center |
| USA 9[16] | *SRR755252, SRR762349*<br>*SRR762350, SRR762351*<br>*SRR764626, SRR764627* | Single (35) | Danold Danforth Center |
| USA 10[17] | SRR1656746 | Single (101) | University of Nebraska - Lincoln |
| USA 11[18] | SRR1567899 | Paired (50) | Iowa State University |
| USA 12*[19] | SRR504480 | Single (100) | University of California - Berkeley |
| USA 13[20] | SRR1587038 | Single (101) | University of Wisconsin - Madison |
| USA 14[21] | SRR1231518 | Single (100) | Cornell University |
| USA 15[22] | SRR1272115 | Paired (50) | DuPont Pioneer |
| USA 16[23] | SRR640263 | Single (35) | Yale University |
| USA 17[24] | *SRR520998, SRR520999* | Paired (51) | Cold Spring Harbor Laboratory |
| USA 18[25] | SRR536834 | Single (76) | Virginia Tech |
| USA 19[26] | SRR999052 | Paired (50) | Cold Spring Harbor Laboratory |
| USA 20[27] | SRR248565 | Paired (81) | Stanford University |
| CHN 1[28] | SRR491307 | Paired (76) | China Agricultural University |
| CHN 2[29] | SRR1522119 | Paired (102) | China Agricultural University |
| CHN 3[30] | SRR910231 | Paired (91) | China Academy of Agricultural Sciences |
| DEU 1[31] | SRR924107 | Single (96) | MPIPZ |
| DEU 2[32] | SRR1030995 | Single (85) | University of Bonn |

\*:USA 12 harbors an long introgression on chromosome 2.

### Alignment and initial SNP calling

Low quality sequences were removed using Trimmomatic-0.33 with settings LEADING:3, TRAILING:3, SLIDINGWINDOW:4:15, MINLEN:36 [33]. Trimmed reads were aligned to the repeat masked version of the maize reference genome (version B73 RefGen v3) [8] downloaded from Ensemble (ftp://ftp.ensemblgenomes.org/pub/plants/release-22/fasta/zea_m_ays/dna/) using GSNAP in version 2014-12-29 (with parameters ‐N1,-n 2,-Q) [34]. Output files were converted from SAM to BAM format, sorted, and indexed using SAMtools [35]. SNPs were called in parallel along ten chromosomes of the maize version 3 using SAMtools mpileup (-I ‐F 0.01) and bcftools call (-mv-Vindels ‐Ob).

### SNP list generation

The view function of Bcftools was combined with in-house Python scripts to extract the content of bcf files and classify SNPs based on the number of reference and non-references alleles on every screened SNP locus. In detail, if the total number of reads covering a particular SNP in a particular sample was below 5, then the site was treated as missing data. When 99% reads on the locus of a sample were from the non-reference allele the sample was coded as homozygous non-reference allele. The same criteria were used for calling a site as homozygous reference allele. When the reads containing reference and non-reference alleles totaled more than 90% of all reads and each allele was represented by more than 20% of aligned reads the site was coded as heterozygous. If two or more alleles were present at >1% of aligned reads but the above criteria were not satisfied, the site was also coded as missing data. To reduce the prevalence of false SNPs resulting from the alignment of reads from multiple paralogous loci to a single position in the reference genome, sites which were scored as heterozygous in more than 20% of all genotyped individuals were discarded. In total, 13,360 SNPs were used in downstream analysis. For each of these SNPs, the impact of the SNP on gene function was estimated using SnpEff v4.1 and SnpEff databases (*AGPv3*.26) [36].

### Population structure analysis

The distribution of the three possible genotypes (homozygous reference allele, homozygous non-referenece allele and heterozygous allele) over each of the ten chromosomes of maize was visualized using matplotlib. PhyML 3.0 [37] was used to construct a phylogenetic tree with 100 bootstrap replicates, and 13,360 SNPs in total of 27 data sets.

### Expression bias test

Individual FPKM (Frequency per kilobase of exon per million reads) value for each gene in each data set was calculated using Cufflinks v2.2.1 [38]. Expression values were averaged across all China and USA South samples (excluded USA 12 sample that contained a unique introgressed region) separately. Only genes with average FPKM values >= 10 in both groups were retained for testing expression bias. The remaining genes were sorted into two groups: genes located in the 7 chromosome intervals where USA South and China showed different haplotypes and genes outside these intervals.

### Origins of haplotype blocks

The origins of haplotype blocks observed in some B73 accessions but not in the published reference genome were investigated using data from diverse maize lines in the HapMap2 project [39]. In order to make comparisons to these data, alignments and SNP calling were performed a second time as above using B73 RefGen v2. All of samples in China or USA North clade were combined to generate a consensus sets of SNP calls with reduced missing data. In examining region c2r2, sample USA 12 was used individually in addition to the combined China and USA North sequences (Additional file 1: Figure S1). In the analysis of region c5r2 (Additional file 1: Figure S1), USA 10, USA 14 and USA 15 were combined to generate a consensus set of SNP calls for the UC-Berkeley clade. The resulting SNP sets were employed for phylogenetic analysis as described above, with the alteration that the an approximate likelihood ratio test (aLRT) method with SH-like was employed. The resulting trees were visualized using FigTree v1.4.2 (http://tree.bio.ed.ac.uk/software/figtree/).

## Results

### Relationship among accessions labeled as B73

After alignment, SNP calling, and filtering (see Methods), a total of 13,360 high confidence segregating SNPs were identified among the 27 RNA-seq samples labeled as B73 employed in this study, substantially lower than the '64,000 high quality SNPs identified by RNA-seq in a population segregating for a single non-B73 haplotype [40]. Phylogenetic analysis identified three distinct clades of samples separated by long branches with 100% bootstrap support (Figure 1). One clade consisted entirely of Chinese samples, one clade of samples from US research groups from Minnesota and Wisconsin, and the final clade encompassed the majority samples from US research groups as well as all German samples and the published reference genome for B73. We designated these clades “China”, “USA North”, and “USA South” respectively. Notably, the USA North clade is paraphyletic with respect to the China clade, suggesting B73 samples in China are likely derived from this group while both German samples are clearly part of the USA South Clade.

**Figure 1.**
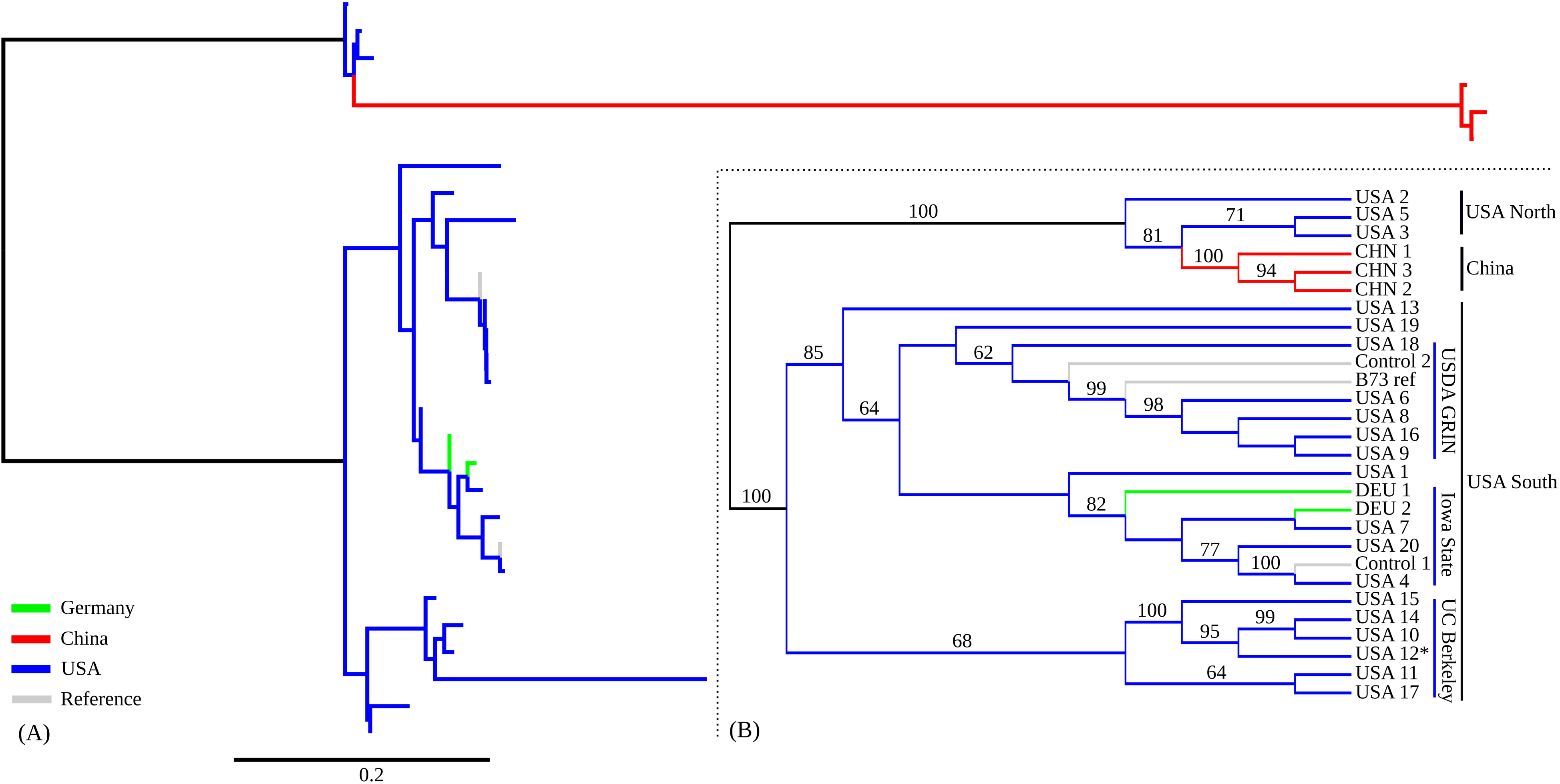
Phylogenetic tree of 27 data sets (A) Distance-scaled branch lengths; (B) Un-scaled tree. Only bootstrap values greater than or equal to 60 are displayed.

The USA South clade was somewhat arbitrarily divided into three subclades with at least 60% bootstrap support, as well as a number of singleton lineages (USA 1, USA 13, USA 19). Two of these clades contained control samples generated for this study, one from B73 seed requested through the USDA Germplasm Resource Network, and one from B73 seed requested from Iowa State. The subclade containing the known USDA B73 sample also contained the B73 reference genome sequence, consistent with the reported seed source for the B73 used in the construction of the reference genome. The final subclade did not contain any control samples. However, it was notable that four of the six samples placed in this clade originated in research groups whose PIs had conducted either PhD or Postdoctoral training with Michael Freeling at UC-Berkeley, and none of the samples outside of this clade originated in research groups linked to UC-Berkeley. Based on these, we designated the final USA South subclade “UC-Berkeley”. This accessions has also been described as “Freeling B73” [41].

### Genomic distribution of within-B73 polymorphisms

The polymorphic SNPs identified in this study could originate from one of several sources including de novo mutations or the introgression of non-B73 haplotypes in one or more lineages. SNPs originating from de novo mutations would be expected to show a distribution approximating that of gene density across the maize chromosomes. SNPs resulting from introgression of other haplotypes into B73 should be tightly clustered.

When the positions of the SNPs identified in this study were plotted it became clear that 55.3% SNPs fall within a small number of dense genomic blocks on chromosomes 2, 3, 4, 5, and 6 (Figure 2). The distribution of non-reference-genome-like haplotype blocks is consistent with the clade relationships identified above. The USA North clade can be defined by a large block of SNPs on chromosome 2, and smaller blocks on chromosomes 2, 3, and 5, all of which are shared with the China B73 clade. In addition to the blocks shared with the USA North B73 clade, samples from the China B73 clade all share a number of additional non-reference-genome-like blocks on chromosomes 2, 4, and 6. There are no non-reference-genome-like blocks shared by all members of the USA South clade, however a single non-reference-genome-like block on chromosome 5 is shared by the UC-Berkeley subclade of USA South. This block appears to share one breakpoint but not both with a block present in the USA North and China samples. The large block non-reference-genome-like block like SNPs observed only on chromosome 2 on USA 12 can likely be explained by the unique origin of this sample from wild type siblings of knotted1 mutants backcrossed into B73 [42]. The remaining USA South samples, including the USDA GRIN, Iowa State, and German samples do not contain any obvious SNP blocks.

**Figure 2.**
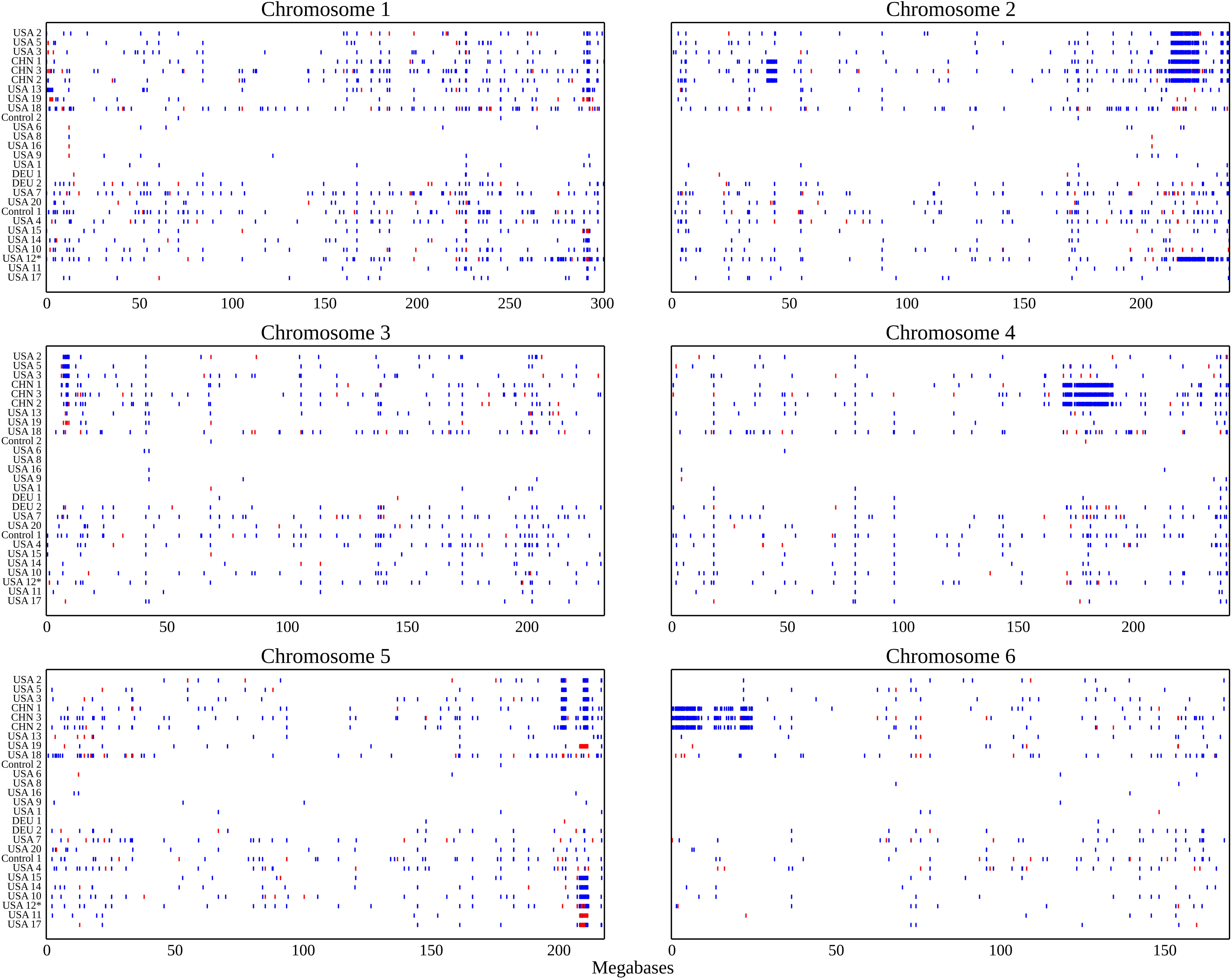
SNP distribution pattern along 6 chromosomes of 27 data sets. Blue dots are non-reference like homozygous alleles and red dots are heterozygous alleles.

### Functional impact of within-B73 polymorphism

Because the data used here came entirely from RNA-seq studies, our ability to detect SNPs was limited to genes which were consistently expressed at high enough levels to provide coverage of target regions. A total of 25,644 genes were expressed at levels >10 FPKM when at least one of data sets analyzed in this study. Of these genes, 633 (2.5%) fell within regions with non-reference-genome-like SNP blocks in one or more B73 clades. Using SnpEff, we identified 10 cases where SNPs produced “high impact” change such as the gain or loss of a stop code or the alteration of a splice donor or splice acceptor site and 396 cases which produced missense mutations which altered protein sequence. Only three genes with reported mutant phenotypes (whp1, mop1, and gol1) were in these regions, which only constituted at 2.7% of 112 classical identified maize genes with reported mutant phenotypes [43]. However, it must be noted that this is likely an underestimate of the true number of changes, nonsense mediated decay may reduce or eliminate the expression of alleles of genes containing high impact SNPs, reducing the chances these SNPs will be detected from RNA-seq data.

### Impact of within-B73 polymorpism on estimated gene expression

The alignment rate for RNA-seq data from non-B73 genotypes to the B73 reference genome is approximately 13% lower than the alignment rate of RNA-seq data generated from B73 plants [44]. To test whether there is a bias towards lower estimated expression levels from RNA-seq data for genes in non-reference-genome-like blocks, the expression of highly expressed genes (ie average expression >=10 FPKM) was compared between samples in the USA South clade (excluding USA 12) and samples in the China clade. Genes within introgressed regions showed a 5.6% reduction on expression in China samples, relative to a control set of genes outside introgressed regions between B73 USA South and B73 China (see Methods). This reduction approximately half as large as would be predicted if the reduced alignment rate of data from non-B73 samples resulted solely from increased difficulty of aligning reads containing SNPs to the reference genome. Potentially, the other half of the reduced alignment rate for non-B73 samples is the result of the expression of lineage specific genes, as previously suggested [44].

### Origins of polymorphic regions in B73 accessions

A total of 7 chromosome intervals (referred to here as c2r1, c2r2, c4r1, c5r1, c5r2, c6r1 and c6r2) containing non-reference genome haplotypes were identified in two or more samples (Table 2; Additional file 1: Figure S1). SNP calls were extracted from individual non-reference-genome-like blocks using the previous version of the maize reference genome (B73 RefGen v2) and compared to genotype calls generated from 103 diverse inbreds resequenced by the Maize HapMap2 project [39]. One example, c2r1 is shown in Figure 4. The non-reference genome haplotype present in this block for the Chinese samples clusters very closely with W22, an older inbred developed in Wisconsin which has also been widely used in the maize genetics research community. Analysis of the other six large haplotype blocks produced longer branch lengths relative to the accessions represented in the Maize HapMap2 dataset (Table 2). However, in each case the haplotypes generated from each clade containing a non-reference-genome-like block clustered together, confirming that these regions did not result from parallel introgressions covering the same regions of the genome. These was also true from c5r2 which was represented in both the USA North and China clades as well as the UC-Berkeley subclade (Additional file 2: Figure S2D). A constraining the c2r2 region to only cover that portion of the genome which contained a block of SNPs in the USA North clade, the China clade and sample USA 12 revealed that USA North and China clustered together while USA 12 was placed at a different location on the tree (Additional file 2: Figure S2A).

**Table 2.** Relationship of Non-Reference-Genome Like SNP Blocks to Haplotypes Surveyed by HapMap2.

| Genomic blocks | Chr | Start (kb) | Stop (kb) | Closest haplotypes | Branch length | Present in |
| --- | --- | --- | --- | --- | --- | --- |
| c2r1 | 2 | 40000 | 44300 | W22 | 0.00000029 | China |
| c2r2 | 2 | 212450 | 224250 | BKN027 | 0.38965362 | China |
|  |  |  |  | BKN027 | 0.38965366 | USA North |
|  |  |  |  | M162W | 0.31485540 | USA 12 |
| c4r1 | 4 | 169650 | 191550 | W22 | 0.62055176 | China |
| c5r1 | 5 | 201200 | 203000 | no single best match | - | China |
|  |  |  |  | no single best match | - | USA North |
| c5r2 | 5 | 209732 | 211540 | no single best match | - | China |
|  |  |  |  | no single best match | - | USA North |
|  |  |  |  | no single best match | - | USA South |
| c6r1 | 6 | 120 | 8800 | CML511 | 0.59243725 | China |
| c6r2 | 6 | 20900 | 24670 | OH7B | 0.08886910 | China |

**Figure 3.**
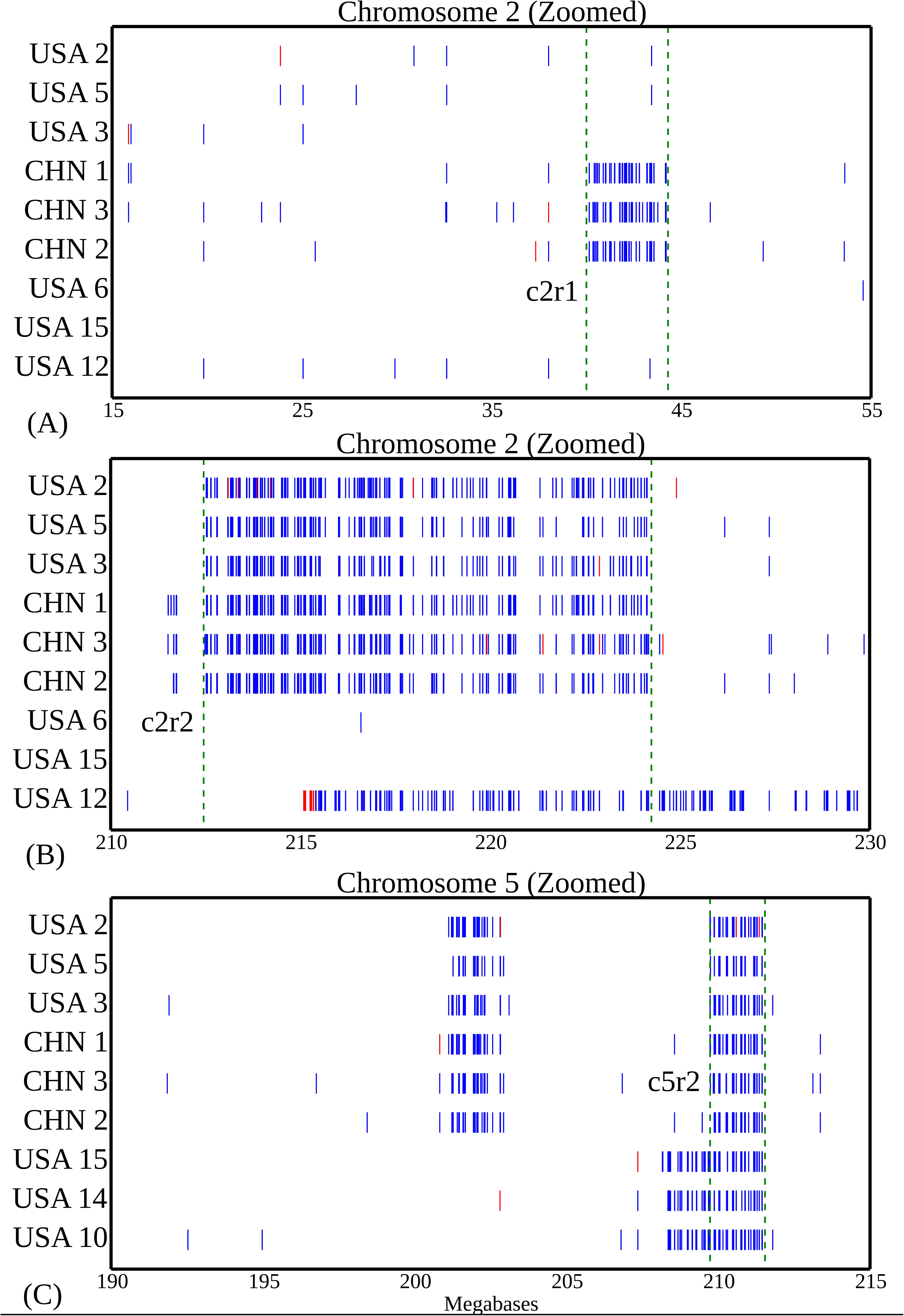
Zoomed haplotype regions of c2r1, c2r2 and c5r2. (A) Selected haplotype region c2r1 on Chromosome 2; (B) Selected haplotype region c2r2 on Chromosome 2; (C) Selected haplotype region c5r2 on Chromosome 5. Blue line is homozygous non-reference allele and red line is heterozygous allele. Regions within green bars are identified haplotype regions.

**Figure 4.**
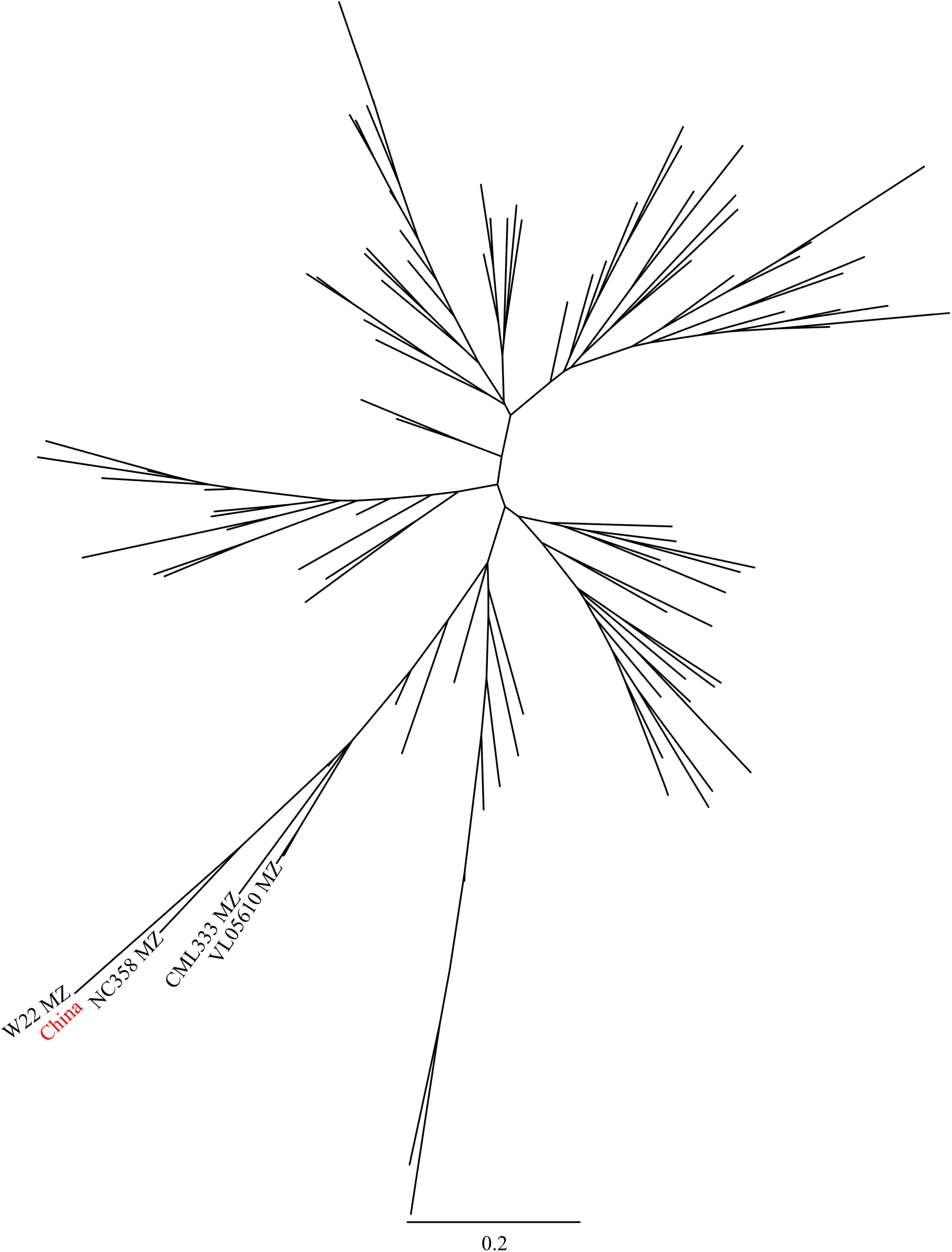
Origin of haplotype region of c2r1.

## Discussion

The maize community has long speculated that significant differences exist among B73 from different sources. Recently that it has become feasible to quantify the specific differences among B73 accessions. Here we employed previously published RNA-seq data sets from a large number of independent research groups to assess the diversity among B73 accessions. No cases of samples which were labeled as originated from B73 but were clearly not B73 based on SNP data were identified in this study. Despite a 40+ generation reproductive history distributed across at least three continents, this analysis shows that 97.7% of the gene space of the maize genome is represented by a single consistent haplotype across all B73 accessions included in this study.

The interspersed SNPs distributed over ten chromosomes of maize may result from de-novo mutations, segregation of heterozygous loci in the original B73 founder accession [4], or false positive SNP calling errors. However, the polymorphic differences identified among B73 samples in this study primarily fell into a small number of dense non-reference-genome-like blocks which would be consistent with introgression of non-B73 germplasm into a B73 background. It is important to note that the B73 reference genome was sequenced recently relative to the total age of the B73 accession. Therefore, it is not possible to infer whether a given non-reference-genome-like block originated from introgression into the line in which the non-reference-genome SNPs are observed or introgression into the B73 lineage which was ultimately employed in the creation of the B73 reference genome. However, in either case the relatively small size of these non-reference genome like blocks suggests multiple generations of backcrossing to the original B73 line, which would not be consistent with a model based on unrecognized pollen contamination.

Instead we propose a model based on the results from Sample USA 12. Sample 12 consists of homozygous wild-type plants selected from family segregating for the Knotted1 [19]. Therefore the block on chromosome 2 (̃1% of the total maize genome) likely represents residual sequence from the knotted1 mutant donor parent line and is consistent with at least 5 generations of backcrossing (expected contribution of the donor parent = ̃1.56%). Similar accidental fixations of unlinked regions may have occurred during the intentional introgression of other traits into a B73 background, such as disease resistance genes [45].

The monophyletic placement of Chinese B73 datasets suggests that the B73 seed available in China likely originated from a single transfer from the USA, apparently of seed belonging to the USA North clade and is an indicator of current tight controls on seed import/export which limit the ease with which seed change be exchanged between collaborators in China and the United States. Samples from Germany did not consistently form a monophyletic group. The concordance of academic lineages and genomic relationships in the UC Berkeley subclade acts as a remarkable positive control. More extensive sampling of B73 samples from many labs which employ this genotype in maize genetics research but have not, to date, published RNA-seq datasets may identify further B73 clades and subclades and additional cases where specific genomic variations have dispersed across the country as graduate students and postdocs leave a given lab for faculty positions of their own.

## Conclusions

The existence of genomic variation among samples labeled as belonging to the same accession creates barriers to reproducibility, one of the core requirements of the scientific method. In this study no examples of sample mislabeling were identified. However, a number of non-reference-genome-like blocks were identified in B73 samples originated from some sources. These blocks were shown to contain missense and nonsense mutations and measurably lower estimated expression values for genes in these regions. The identification of the relationships among different variants of B73 and the genomic locatons of non-reference-genome-like regions will allow these differences to be controlled for future studies. With the rapid rise of sequencing-based assays such as RNA-seq, the strategy employed here may be a good one to apply in any case where one or more reference genotypes are widely employed in research across institutions, countries, and continents.

## Author’s contributions

JCS and ZL designed the study, ZL collected the data, ZL performed the analysis, and JCS and ZL wrote the manuscript.

## Acknowledgements

This project is supported by start-up funds from University of Nebraska-Lincoln to JCS.

## Additional Files

Additional file 1: Figure S1. Identified 7 haplotype blocks and SNP distribution pattern along 10 chromosomes of 27 data sets.

Additional file 2: Figure S2. Phylogenetic tree of other haplotype regions and corresponding lines in HapmapV2.(Additional file 2:Figure S2A, Origin of c2r2; Additional file 2:Figure S2B, Origin of c4r1; Additional file 2:Figure S2C, Origin of c5r1; Additional file 2:Figure S2D, Origin of c5r2; Additional file 2:Figure S2E, Origin of c6r1; Additional file 2:Figure S2F, Origin of c6r2.)

